# LanA (Language Atlas): A probabilistic atlas for the language network based on fMRI data from >800 individuals

**DOI:** 10.1101/2022.03.06.483177

**Authors:** Benjamin Lipkin, Greta Tuckute, Josef Affourtit, Hannah Small, Zachary Mineroff, Hope Kean, Olessia Jouravlev, Lara Rakocevic, Brianna Pritchett, Matthew Siegelman, Caitlyn Hoeflin, Alvincé Pongos, Idan A. Blank, Melissa Kline Struhl, Anna Ivanova, Steven Shannon, Aalok Sathe, Malte Hoffmann, Alfonso Nieto-Castañón, Evelina Fedorenko

**Affiliations:** Massachusetts Institute of Technology; Johns Hopkins University; Carnegie Mellon University; Carleton University; Columbia University; University of Chicago; University of California, Berkeley; University of California, Los Angeles; Harvard, Mass General Hospital; Boston University

## Abstract

Two analytic traditions characterize fMRI language research. One relies on averaging activations voxel-wise across individuals. This approach has limitations: because of inter-individual variability in the locations of language areas, a location in a common brain space cannot be meaningfully linked to function. An alternative approach relies on identifying language areas in each individual using a functional ‘localizer’. Because of its greater sensitivity, functional resolution, and interpretability, functional localization is gaining popularity, but it is not always feasible, and cannot be applied retroactively to past studies. We provide a solution for bridging these currently disjoint approaches in the form of a *probabilistic functional atlas* created from fMRI data for an extensively validated language localizer in 806 individuals. This atlas enables estimating the probability that any given location in a common brain space belongs to the language network, and thus can help interpret group-level peaks and meta-analyses of such peaks, and lesion locations in patient investigations. More meaningful comparisons of findings across studies should increase robustness and replicability in language research.

## Background and Summary

fMRI is an invaluable non-invasive tool for illuminating the brain’s architecture, especially for human-unique abilities like language. A common analytic approach in fMRI language studies is to average activation maps voxel-wise in a common brain space and perform statistical inference across individuals in each voxel. However, because of the well-established inter-individual variability in the locations of functional areas in the association cortex (Frost &Goebel, 2012; Tahmasebi et al., 2012), activations do not line up well across individuals, leading to low sensitivity and functional resolution (Nieto-Castañón &Fedorenko, 2012). Further, the results of group-averaging analyses are generally interpreted through reverse inference from anatomical locations to function (Fedorenko, 2021; Poldrack, 2011), but because of the variability mentioned above, combined with the functional heterogeneity of the association cortex, locations in a common brain space cannot be meaningfully linked to function (see Fedorenko &Blank, 2020, for a discussion of this issue for ‘Broca’s area’).

An alternative analytic approach, which circumvents voxel-wise brain averaging, is known as ‘functional localization’ (Nieto-Castañón &Fedorenko, 2012; Saxe, 2006). In this approach, a brain region or network that supports a mental process of interest is identified with a functional contrast in each individual and then its functional responses to some new critical condition(s) are examined. This approach yields greater sensitivity, functional resolution, and interpretability, and has been highly successful across many domains of perception and cognition, including language. As a result, many research groups are now moving away from group-averaging analyses toward individual-subject analyses (e.g., Fedorenko, 2021; Gratton &Braga, 2021).

However, functional localization is not always feasible. Further, although studies that rely on functional localization can be straightforwardly compared to each other, it is at present unclear how to relate the results from such studies to group-averaging fMRI studies, or other studies that rely on brain averaging (e.g., studies that use voxel-based morphometry (VBM) or voxel-based lesion-symptom mapping (VLSM) in patient work; Wilson, 2017). To help bridge the gap between these two different analytic traditions in language research, we created a ***probabilistic functional atlas of the language network (‘Language Atlas’ or LanA)*** by overlaying 806 individual activation maps for a robust and validated language ‘localizer’ (Fedorenko et al., 2010; Mahowald &Fedorenko, 2016). This localizer relies on a contrast between the processing of sentences and a linguistically/acoustically degraded condition and is robust to changes in materials, modality of presentation, and task (see <u>Methods</u>). This localizer identifies the fronto-temporal language network that selectively (Fedorenko et al., 2011) supports high-level language comprehension and production, including the processing of word meanings and combinatorial syntactic/semantic processing (Bautista &Wilson, 2016; Fedorenko et al., 2012, 2020). Further, a network that closely corresponds to this functional contrast emerges from task-free resting state data (Braga et al., 2020).

LanA allows one to estimate for any location in a common brain space the probability that it falls within the language network. In this way, this atlas can provide a common reference frame and help interpret a) group-level activation peaks from past and future fMRI studies, b) results of meta-analyses of such peaks (Hauptman et al., 2022), c) lesion locations in individual brains (Woolgar et al., 2018) or lesion overlap loci in VBM/VLSM analyses, d) electrode locations in ECoG/SEEG investigations and locations of source-localized activity in MEG studies. Furthermore, LanA e) can help guide functional mapping during brain surgery when fMRI is not possible, f) can be related voxel-by-voxel to any whole-brain data (Markello et al., 2022), including structural data, gene expression data (Richiardi et al., 2015), or receptor density data (Hansen et al., 2021), in order to ask whether/how these features correlate with the language network’s topography, and g) can help select patches in post-mortem brains for cellular analyses to maximize the chances of examining language cortex. (We emphasize that LanA is not a replacement for localizers: when possible, a localizer task should be performed. As we show in **SI-1**, the effect sizes obtained from group-level ROIs based on LanA, or from commonly used Glasser parcels (Glasser et al., 2016) are underestimated.)

We make the atlas available for two most commonly used brain templates (**Figure 1**): a volume-based MNI template (IXI549Space; SPM12; Friston, 2007) and a surface-based FSaverage template (Fischl et al., 1999). We also release i) individual activation maps (in the MNI and FSaverage spaces), along with demographic data, and ii) individual-level neural markers (based on the volumetric analyses), including effect sizes, activation extent, and stability of activation across runs. The neural marker data can be used as normative distributions against which any new population (e.g., children or individuals with developmental or acquired brain disorders) can be evaluated.

**Figure 1:**
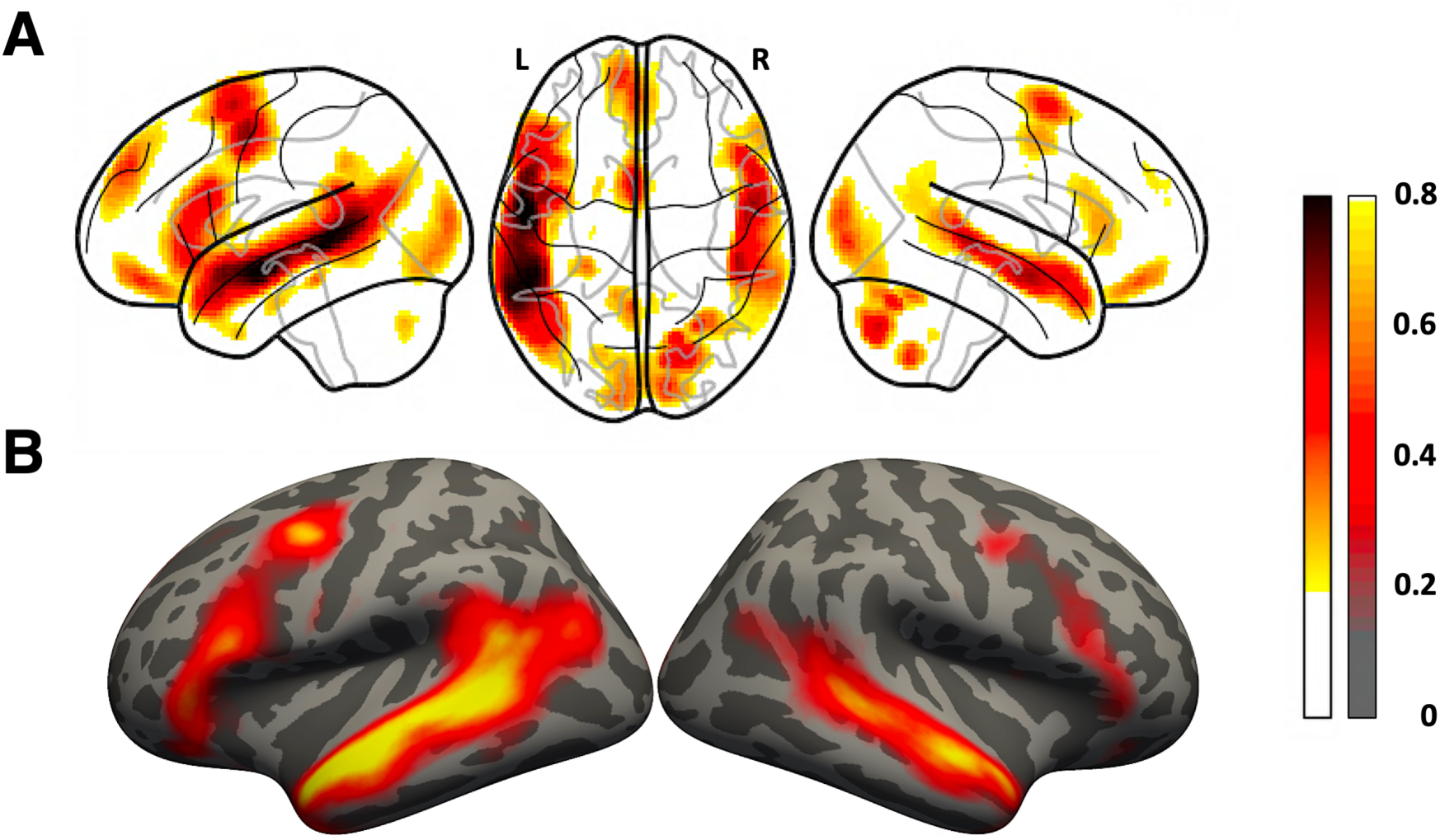
Probabilistic functional atlas for the *language > control* contrast based on overlaid individual binarized activation maps (where in each map, the top 10% of voxels are selected, as described in the text). *A)* SPM-analyzed volume data in the MNI template space (based on 806 individual maps). *B)* FreeSurfer-analyzed surface data in the FSaverage template space (based on 804 individual maps). In both figures, the color scale reflects the proportion of participants for whom that voxel belongs to the top 10% of *language > control* voxels.

## Methods

### Participants

A total of 806 neurotypical adults (477, ∼59%, female), aged 19 to 75 (average: 30.23; standard deviation: 7.08; median: 29), participated for payment between September 2007 and June 2021, as summarized in **Table 1**. All participants had normal or corrected-to-normal vision, and no history of neurological, developmental, or language impairments. Handedness information was collected for 758 (∼94%) of the 806 participants. Of those, 707 participants (∼93%) were right-handed, as determined by the Edinburgh handedness inventory (Oldfield, 1971) or self-report, 38 (∼5%) were left-handed, and 13 (∼2%) – ambidextrous. (The participants for whom handedness is missing in the database are most likely right-handed because most of them were tested during the earlier years of data collection when right-handedness was one of the requirements for participation.) Of the 806 participants, 629 (∼78%) were native speakers of English, and the remaining 177 (∼22%) – native speakers of another language and proficient speakers of English (see Malik-Moraleda, Ayyash et al., 2021, for evidence that the topography of language responses for a language that an individual is proficient in is similar to that of their native language, and see **SI-2** for a comparison between the atlas generated using all 806 participants, vs. only the 629 native English speakers).

**Table 1:** Summary demographics of the 806 participants included in the atlas.

| Summary Demographics of this Population |  |  |
| --- | --- | --- |
| Age | 30.23 ± 7.08 | Years |
| Gender | 40.57% | Male |
|  | 59.18% | Female |
| Handedness | 87.72% | Right-Handed |
|  | 4.71% | Left-Handed |
|  | 1.61% | Ambidextrous |
|  | 5.96% | No Handedness Info |
| Native English Speaker Status | 78.04% | Native Speakers of English |
|  | 21.96% | Proficient Speakers of English |

Each participant completed a language ‘localizer’ task (Fedorenko et al., 2010) as part of one of the studies in the Fedorenko lab. Each scanning session lasted between 1 and 2 hours and included a variety of additional tasks. All participants gave informed written consent in accordance with the requirements of the MIT’s Committee on the Use of Humans as Experimental Subjects (COUHES).

### Participant and session selection

The 806 scanning sessions above (one session per participant) were selected from a total of 1,065 sessions across 819 participants that were available in the Fedorenko Lab database as of June 2021. The goal was to include as many participants as possible and, for the 163 participants who performed a language localizer in multiple sessions, to select a single session with high-quality data. To assess data quality, we examined the stability of the activation topography for the language localizer contrast (see <u>Language localizer paradigm</u>) across runs. This analysis was performed on the data preprocessed and analyzed in the volume (i.e., SPM-based analyses; see <u>SPM preprocessing and analysis pipeline</u>). For 1,062 of the 1,065 sessions, we calculated voxel-wise spatial correlations in activation magnitudes the *language > control* contrast (see <u>Language localizer paradigm</u>) between the odd-numbered and even-numbered runs (three remaining sessions consisted of a single run and were evaluated by visual inspection of the contrast maps). The correlation values were calculated within the language ‘parcels’—masks that denote typical locations of language areas. These masks (available at http://evlab.mit.edu/funcloc) were derived from a probabilistic language atlas based on 220 participants (a subset of the participants in the current set of 806) and have been used in much past work (e.g., Diachek, Blank, Siegelman et al., 2020; Ivanova et al., 2020; Jouravlev et al., 2019, 2020; Mollica et al., 2020; Shain, Blank et al., 2021; Wehbe et al., 2021). Six masks (three in the frontal cortex and three in the temporal and parietal cortex) were derived from the probabilistic atlas in the left hemisphere and mirror-projected onto the right hemisphere. For each session, the correlation values were averaged across the twelve parcels, leading to a single value per session. This spatial correlation measure quantifies the stability of the activation landscape and is an objective proxy for data quality; it is affected by factors like head motion or sleepiness, but does not require subjective visual inspection of contrast maps. Sessions where the spatial correlation value was negative (n=23; ∼2%) were excluded, leaving 1,042 sessions across 806 participants. For the 163 participants with more than one session, we selected the session with the highest spatial correlation value for the inclusion in the atlas (see Lipkin et al., In prep. for evidence of the stability of spatial correlation values across sessions: i.e., if a participant shows a high spatial correlation in one session, they are likely to show a high spatial correlation in another session). Following this data selection procedure, the Fisher-transformed spatial correlations of the participants’ *language > control* contrast were r=0.98 and r=0.57 for the left and right hemispheres, respectively (see Mahowald &Fedorenko, 2016, for similar values on a subset (n=150) of these participants).

### Language localizer paradigm

Across the 806 participants, ten language localizer versions were used, as summarized in **Table 2**. In each version, a sentence comprehension condition was contrasted with a linguistically or acoustically degraded control condition. Visual (reading) and auditory (listening) contrasts have been previously established to engage the same fronto-temporal language network (e.g., Chen et al., 2021; Fedorenko et al., 2010; Malik-Moraleda, Ayyash et al., 2021; Scott et al., 2017). Activity in this network has further been shown to not depend on task or materials (Fedorenko et al., 2010) and to show robust effects across typologically diverse languages (Malik-Moraleda, Ayyash et al., 2021). Furthermore, this network can be recovered from naturalistic task-free (resting state) data based on patterns of BOLD signal fluctuations over time (Blank et al., 2014; Braga et al., 2020), and corresponds nearly perfectly to the network based on the *sentences > nonwords* contrast (Fedorenko et al., 2010). As a result, we pooled data from across the different versions in the current study (see **SI-3** for a reality-check analysis showing robust *language > control* effects across all ten versions, and **SI-2** for evidence that an atlas defined on only ***Localizer Version A*** is nearly identical to that which leverages data from all versions).

**Table 2:** Timing parameters for each version of the language localizer task. Under task type, the options are defined as follows: BP = Button Press, MP = Memory Probe, N = No Task.

| Version | A | B | C | D | E |
| --- | --- | --- | --- | --- | --- |
| Number of participants | 624 | 67 | 60 | 15 | 17 |
| Task type | BP | MP | N/MP | MP | MP |
| Words / Nonwords per trial | 12 | 8 | 12 | 12 | 8 |
| Modality | Visual | Visual | Visual | Visual | Visual |
| Trial duration (ms) | 6000 | 4800 | 6000 | 6000 | 4800 |
| Fixation | 100 | 300 | 300 | 300 | 300 |
| Stimulus | 450 | 350 | 350 | 350 | 350 |
| Button icon / Memory probe | 400 | 1350 | 0 or 1000 | 1000 | 1350 |
| Fixation | 100 | 350 | 500 or 1500 | 500 | 350 |
| Trials per block | 3 | 5 | 3 | 3 | 5 |
| Block duration (s) | 18 | 24 | 18 | 18 | 24 |
| Blocks per condition per run | 8 | 4 | 8 | 6 | 4 |
| Conditions | Sentences | Sentences | Sentences | Sentences | Sentences |
|  | - | Wordlists | - | Wordlists | Wordlists |
|  | - | - | - | - | Jabberwocky |
|  | Nonwords | Nonwords | Nonwords | Nonwords | Nonwords |
| Fixation block duration (s) | 14 | 16 | 18 | 18 | 16 |
| Number of fixation blocks | 5 | 3 | 5 | 4 | 5 |
| Total run time (s) | 358 | 336 | 378 | 396 | 464 |
| Number of runs | 2 | 2-5 | 2 | 2-3 | 6-8 |

| Version | F | G | H | I | J |
| --- | --- | --- | --- | --- | --- |
| Number of participants | 6 | 4 | 8 | 4 | 1 |
| Task type | BP | MP | N | MP | N |
| Words / Nonwords per trial | 12 | 8 | Variable | 8 | Variable |
| Modality | Visual | Visual | Auditory | Auditory | Auditory |
| Trial duration (ms) | 4800 | 4800 | 18000 | 6000 | 12000 |
| Fixation | 600 | 300 |  |  |  |
| Stimulus | 350 | 350 |  | 3300-4300 |  |
| Button icon / Memory probe |  | 1350 |  | 1000 |  |
| Fixation |  | 350 |  | Until 6000 |  |
| Trials per block | 5 | 5 | 1 | 4 | 1 |
| Block duration (s) | 24 | 24 | 18 | 24 | 12 |
| Blocks per condition per run | 4 | 8 | 8 | 4 | 16 |
| Conditions | Sentences | Sentences | Intact | Sentences | Sentences |
|  | Wordlists | - | - | Wordlists | - |
|  | Jabberwocky | - | - | Jabberwocky | - |
|  | Nonwords | Nonwords | Degraded | Nonwords | Nonwords |
| Fixation block duration (s) | 25 | 16 | 14 | 16 | 16 |
| Number of fixation blocks | 4 | 5 | 5 | 5 | 5 |
| Total run time (s) | 504 | 464 | 358 | 464 | 464 |
| Number of runs | 2-8 | 2 | 2 | 4 | 2 |

The vast majority of participants (624, ∼77.4%) performed ***Localizer version A*** – a reading version, where sentences and nonword strings are presented one word/nonword at a time at the rate of 450ms per word/nonword, in a blocked design (with 3 sentences/nonword strings in each 18s block). Participants were instructed to read attentively and to press a button at the end of each trial, when a picture of a finger pressing a button appeared on the screen. The experiment consisted of two ∼6-minute-long runs, for a total of 16 blocks for each of the two conditions. The presentation script and stimuli for this localizer version can be downloaded at http://evlab.mit.edu/funcloc/ (for the stimuli used in the other localizer versions, contact EF). ***Localizer versions B-G*** (performed by 169 participants, ∼21.0%) also used visual presentation and the same contrast as in version A, with each word/nonword presented at the rate of 350ms per word/nonword. Participants were instructed to either read attentively (with no additional task), read attentively and press a button at the end of each trial, or read attentively and perform a memory probe task at the end of each trial (deciding whether a word/nonword appeared in the string just read). Versions B, D, E, and F included additional conditions besides the critical sentences and nonword-strings conditions, but we just focus on the latter two conditions here. Each block in versions B-G consisted of either 3 or 5 sentences/nonword strings and lasted 18 or 24 seconds. The experiments consisted of between 2 and 8 runs (for a total of 8-32 blocks per condition), with the runs lasting between ∼5.5 and ∼8.5 minutes. Finally, ***Localizer versions H-J*** (performed by 13 participants, ∼1.6%) used auditory presentation. Participants were instructed to either listen attentively (with no additional task) or listen attentively and perform a memory probe task at the end of each trial. Versions I and J used nonword strings as the control condition (like the visual versions A-G; and version I included additional conditions besides the critical conditions), and version H used an acoustically degraded control condition (see Scott et al., 2017 for details). Each block consisted of several sentences (version I) or a short text (versions H, J) and lasted 12-24 seconds. Experiments consisted of 2 or 4 runs (for a total of 16-32 blocks per condition), with the runs lasting between ∼6 and ∼8 minutes.

### fMRI data acquisition

Structural and functional data were collected on the whole-body, 3 Tesla, Siemens Trio scanner with a 32-channel head coil, at the Athinoula A. Martinos Imaging Center at the McGovern Institute for Brain Research at MIT. T1-weighted structural images were collected in 176 sagittal slices with 1 mm isotropic voxels (TR = 2,530 ms, TE = 3.48 ms). Functional, blood oxygenation level dependent (BOLD), data were acquired using an EPI sequence (with a 90 degree flip angle and using GRAPPA with an acceleration factor of 2), with the following acquisition parameters: thirty-one 4 mm thick near-axial slices acquired in the interleaved order (with 10% distance factor), 2.1 mm x 2.1 mm in-plane resolution, FoV in the phase encoding (A >> P) direction 200 mm and matrix size 96 mm x 96 mm, TR = 2,000 ms and TE = 30 ms. Prospective acquisition correction (Thesen et al., 2000) was used to adjust the positions of the gradients based on the participant’s motion from the previous TR. The first 10 s of each run were excluded to allow for steady state magnetization.

### SPM preprocessing and analysis pipeline

For the SPM analysis (**Figure 2**), fMRI data were analyzed using SPM12 (release 7487), CONN EvLab module (release 19b) and other custom MATLAB scripts. Each participant’s functional and structural data were converted from DICOM to NIFTI format. All functional scans were coregistered and resampled using B-spline interpolation to the first scan of the first session (Friston et al., 1995). Potential outlier scans were identified from the resulting subject-motion estimates as well as from BOLD signal indicators using default thresholds in CONN preprocessing pipeline (5 standard deviations above the mean in global BOLD signal change, or framewise displacement values above 0.9 mm, Nieto-Castanon, 2020). Functional and structural data were independently normalized into a common space (the Montreal Neurological Institute [MNI] template; IXI549Space) using SPM12 unified segmentation and normalization procedure (Ashburner &Friston, 2005) with a reference functional image computed as the mean functional data after realignment across all timepoints omitting outlier scans. The output data were resampled to a common bounding box between MNI-space coordinates (−90, -126, -72) and (90, 90, 108), using 2mm isotropic voxels and 4^th^ order spline interpolation for the functional data, and 1mm isotropic voxels and trilinear interpolation for the structural data. Last, the functional data were then smoothed spatially using spatial convolution with a 4 mm FWHM Gaussian kernel. Effects were estimated using a General Linear Model (GLM) in which each experimental condition was modeled with a boxcar function convolved with the canonical hemodynamic response function (HRF) (fixation was modeled implicitly). Temporal autocorrelations in the BOLD signal timeseries were accounted for by a combination of high-pass filtering with a 128 seconds cutoff, and whitening using an AR(0.2) model (first-order autoregressive model linearized around the coefficient a=0.2) to approximate the observed covariance of the functional data in the context of Restricted Maximum Likelihood estimation (ReML). In addition to main condition effects, other model parameters in the GLM design included first-order temporal derivatives for each condition, modeling spatial variability in the HRF delays, as well as nuisance regressors controlling for the effect of slow linear drifts, subject-motion parameters, and potential outlier scans on the BOLD signal.

**Figure 2:**
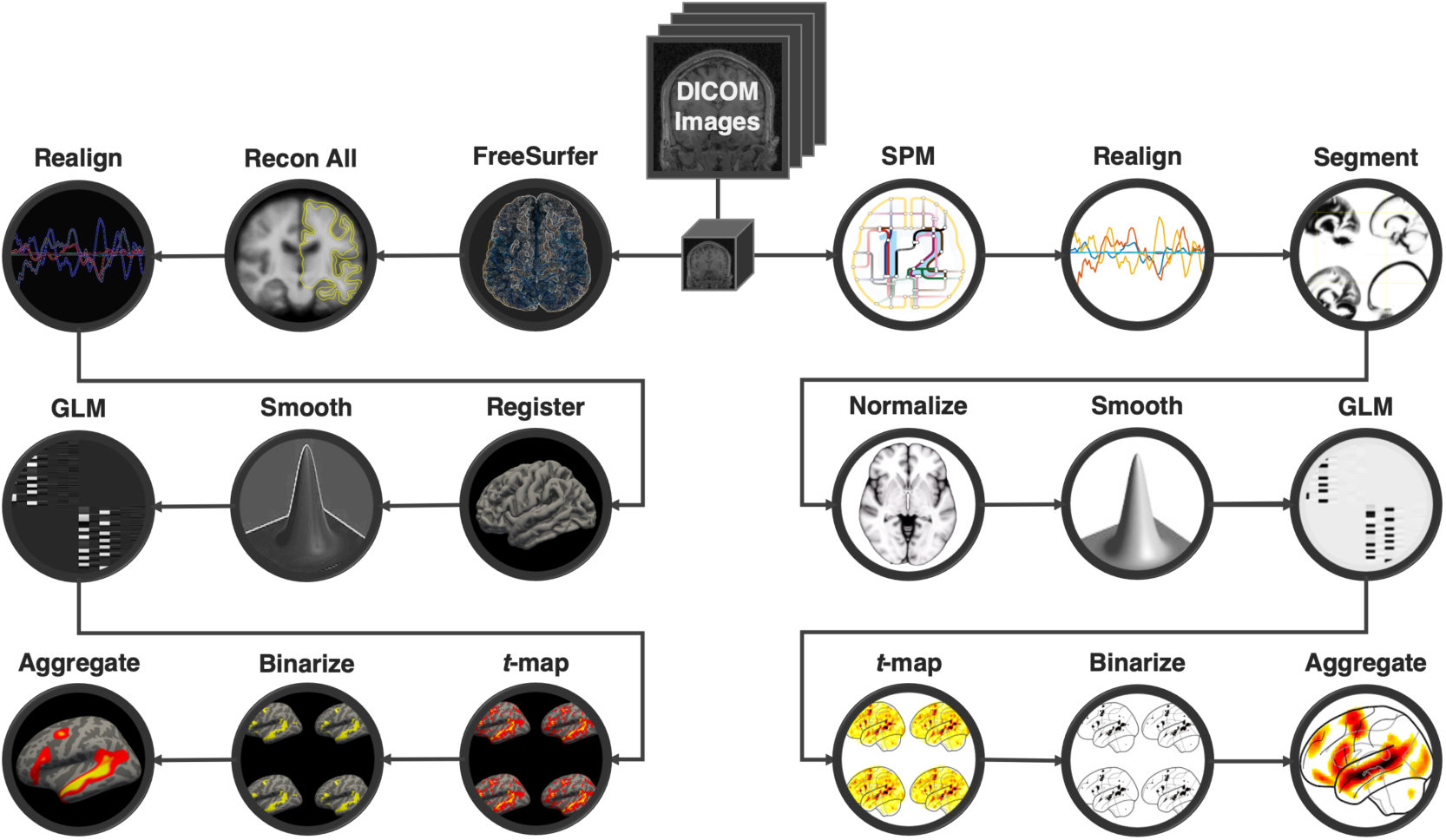
Overview of SPM and FreeSurfer preprocessing and analysis pipelines. Raw dicom images are converted to NIfTI format, motion-corrected, mapped to a common space and smoothed during preprocessing. Each session is then modeled, *t*-maps are extracted and thresholded, and all sessions are aggregated to create the probabilistic atlas.

### FreeSurfer preprocessing and analysis pipeline

For the FreeSurfer analysis (**Figure 2**), fMRI data was analyzed using FreeSurfer v6.0.0. Each participant’s functional and structural data were converted from DICOM to NIFTI format using the default *unpacksdcmdir* parameters. (Two of the 806 participants could not be included in this pipeline because their raw dicom files were lost, leaving 804 participants for this analysis.) The raw data were then sampled onto both hemispheres of the FSaverage surface, motion corrected and registered using the middle time point of each run. The data were then smoothed spatially with a 4 mm FWHM Gaussian filter. Effects were estimated using a GLM in which each condition was modeled with a first order polynomial regressor fitting the canonical HRF (the first 4 time points of each run were excluded to allow for steady state magnetization). The GLM also included nuisance regressors for offline-estimated motion parameters.

### Atlas creation

*SPM:* Using custom code (available at OSF: https://osf.io/kzwbh/), we computed the overlap of the individual activation maps for the *language > control* contrast using the 806 participants analyzed in the SPM12 pipeline. In particular, we used whole-brain *t*-maps that are generated by the first-level analysis and that contain a *t*-value for the relevant contrast in each voxel (a post-hoc analysis compared the whole-brain *t*-maps to their respective unscaled contrast maps and found strong voxel-wise correlations over the set of 806 participants: *r*=0.93±0.03) (see **SI-2** for evidence that atlases generated from *t*-maps vs. contrast maps are highly similar). In each individual map, we selected the 10% of voxels across the brain with the highest *t*-values for the *language > control* contrast (average *minimum t-*value across participants was 1.73 (median: 1.62); average *maximum t*-value was 13.8 (median: 13.7)). We chose the top 10% approach over an approach where each individual map is thresholded at a fixed *t*-value (as in Fedorenko et al., 2010) to account for inter-individual variability in the overall strength of BOLD signal responses due to trait or state factors (Erdoğan et al., 2016; He et al., 2010; Power et al., 2015; Wong et al., 2013) (but see **SI-2** for evidence that an atlas based on the fixed *t*-value thresholding approach yields a nearly identical topography). These maps were then binarized so that the selected voxels got assigned a value of 1 and the remaining voxels—a value of 0. Finally, these values were averaged across participants in each voxel. The resulting atlas contains in each voxel a value between 0 and 1, which corresponds to the proportion of the 806 participants for whom that voxel falls in the 10% of voxels across the brain with the highest *t*-values for the *language > control* contrast. In the left hemisphere, these values range from 0 to 0.82, and in the RH—from 0 to 0.64 (the values are lower in the RH presumably because the majority of the selected voxels fall in the LH: average and median proportions of selected voxels falling in the LH are 58.3% and 57.8%, respectively).

#### FreeSurfer

Using custom code (available at OSF: https://osf.io/kzwbh/), we computed the overlap of the individual activation maps for the *language > control* contrast, using the 804 participants analyzed in the FreeSurfer pipeline. The procedure was similar to that used for the SPM-based atlas, except that the selection of the highest *t-*values was performed on the surface vertices. To maintain hemispheric asymmetries, rather than evaluating each hemisphere separately, as is generally common for FreeSurfer analyses, the top 10% of vertices were selected from the vertices pooled across the LH and RH, as for the SPM-based atlas. For this atlas, in the left hemisphere, the proportion values range from 0 to 0.90, and in the RH—from 0 to 0.80 (these values are expectedly higher than those in the SPM-based atlas given the superiority of surface-based inter-individual alignment; e.g., Fischl et al., 2008).

### Neural Markers

In addition to the population-level atlases, we also provide a set of individual-level neural markers (based on the volumetric SPM analyses) for the language network in each participant. These neural markers include: effect size, voxel count, and spatial correlation. All of these markers have all been shown to be reliable within individuals over time, including across scanning sessions (Lipkin et al., In prep.; Mahowald &Fedorenko, 2016). We provide each of these measures for each of the ROIs constrained by the previously defined language ‘parcels’ (available at http://evlab.mit.edu/funcloc), which include in each hemisphere three frontal parcels (IFG, IFGorb, MFG) and three temporal/parietal ones (AntTemp, PostTemp, AngG), for a total of 12 parcels.

Effect Size was operationalized as the magnitude (% BOLD signal change) of the critical *language > control* contrast. Within each parcel, we defined—for each participant—a fROI by selecting 10% of the mask’s total voxels with the highest *t-values* for the *language* > *control* contrast using all but one run of the data. We then extracted from the left-out run the responses to the language and control conditions and computed the *language > control* difference. This procedure was repeated across all run partitions. This across-runs cross-validation procedure (Nieto-Castañón &Fedorenko, 2012) ensures independence between the data used to define the fROIs and estimate their responses (Kriegeskorte, 2011). In the final step, the estimates were averaged across the cross-validation folds to derive a single value per participant per fROI. Voxel count (extent of activation) was defined as the number of significant voxels for the critical *language > control* contrast at a fixed statistical threshold (p<0.001 uncorrected threshold). Spatial correlation (stability of the activation landscape) was defined—for the voxels falling within the language parcels—as the Fischer-transformed Pearson correlation between the voxel responses for the *language > control* contrast across odd and even runs. As noted above, for all three measures, we provide 14 values per participant: one for each of the 12 ROIs (6 in each hemisphere), and two additional per-hemisphere values (averaging across the 6 ROIs in each hemisphere). See **Table 3** for a summary of these neural markers within the atlas population.

**Table 3:** Summary neural markers for the *language > control* contrast of the 803 participants included in the atlas for whom we have 2 or more runs. Effect sizes reflect the % BOLD Signal Change of the target contrast in the language parcels using an out-of-sample localizer, Significant voxels are defined at p<0.001 uncorrected, and Spatial Correlation is defined as the Fischer-transformed correlation coefficient over the language parcels between the odd and even runs as a marker of stability. LH=Left-Hemisphere; RH=Right-Hemisphere.

| Summary Markers of this Population | Minimum | 0.25 | Median | 0.75 | Max |
| --- | --- | --- | --- | --- | --- |
| Mean LH Effect Size | -0.28 | 0.88 | 1.28 | 1.65 | 3.68 |
| Mean RH Effect Size | -0.40 | 0.20 | 0.43 | 0.73 | 2.51 |
| Number of Significant LH Voxels | 0 | 1197 | 1908 | 2594 | 4887 |
| Number of Significant RH Voxels | 0 | 196 | 495 | 1044 | 4473 |
| Mean LH Spatial Correlation | -0.01 | 0.74 | 1.00 | 1.23 | 1.90 |
| Mean RH Spatial Correlation | -0.29 | 0.33 | 0.57 | 0.78 | 1.75 |

## Data records

The SPM and FreeSurfer atlases are available for download [*website*] (the website will go live upon manuscript acceptance; in the meantime, the atlas is available from BL). Along with the atlases, we make available i) individual contrast and significance maps (for both the volume-based SPM and the surface-based FreeSurfer pipelines; because we had not obtained consent for raw data release, we cannot make publicly available the raw dicom/NIfTI images), and ii) a dataset of individual neural markers, which can be explored with respect to demographic variables or serve as normative distributions against which any new population can be evaluated.

The complete dataset can be accessed at [*website*] via the prepackaged download links. The ‘Download SPM Atlas’ and ‘Download FS Atlas’ links provide a copy of the language atlas in their respective formats. The SPM atlas is a single volumetric NIfTI file, whereas the FS atlas is comprised of two overlay NIfTI files, one for each hemisphere. Under ‘Download All SPM Data’ and ‘Download All FS Data’, each of the individual participant’s data can be downloaded. In particular, for each of the 806 participants (804 for FS), we provide a ‘Demograhics_&_Summary.txt’ file, which contains relevant information as in **Tables 2** and **3**, the individual contrast and significance maps, and a visualization of their individual activation profile in the selected template space.

As well as allowing the user to download the data, the website offers ample opportunities for online exploration and the retrieval of relevant subsets of data. In particular, individual activation maps can be explored under the ‘Explore Activation Maps’ tab, and relevant neural markers can be explored under the ‘Explore Neural Markers’ tab. In addition, data can be filtered by demographic variables including, age, gender, handedness, native English speaker status, language network lateralization, etc., and these subsets can be downloaded, or their maps / neural markers can be explored. This flexible tool allows individual users to access relevant data for their needs without the requirement for offline filtering.

Finally, we provide a version of the language localizer experiment (*Localizer Version A*, which is used for the majority of the participants) for download, as well as some informational videos and references about the language network.

## Technical Validation

The individual participants’ data quality check was performed as described in the <u>Participants and session selection</u> section.

Individual localizer versions were evaluated to confirm they each elicited a strong *language > control* effect, as described in **SI-2**.

The atlas creation process was evaluated with respect to several hyperparameter choices, and remained robust to each decision, including the inclusion of non-native but fluent English speakers, localizer versions, the use of whole-brain maps based on *t-*values vs. contrast values, and definition of the language system as the top 10% of language>control voxels vs. as language>control voxels that pass a specific significance threshold. We outline the minimal impact of all these choices and the strong correlations to the main atlas in **SI-3**.

Finally, in **SI-1**, we demonstrate that group-level ROIs defined based on the highest-overlap voxels in LanA outperform commonly used Glasser ROIs derived from multi-modal Human Connectome Project data (Glasser et al., 2016) in effect size estimation. The latter grossly underestimate effect sizes, especially for the frontal language areas. Of course, as expected (e.g., Nieto-Castañón &Fedorenko, 2012), individual-level language fROIs are still the best for accurately estimating effect sizes, and these outperform the group-based LanA fROIs, but in cases where individual localization may not be possible (e.g., in retroactively re-analyzing past studies), LanA-based group ROIs are recommended, as they fare substantially better than Glasser ROIs.

## Usage Notes

The data records presented in this paper, including materials for download and exploration at the [*website*] are available for free and fair use to individual and academic researchers, institutions, and entities provided that this work is appropriately referenced. Although this atlas has potential for clinical applications, the authors assume no responsibility for the use or misuse of LanA and associated data records in clinical and other settings.

## Code availability

Code associated with this manuscript can be found at OSF: https://osf.io/kzwbh/.

## Acknowledgements

EF and some of this work were supported by NIH awards K99R00 HD057522, R01 DC016607 (and supplement 3R01DC016607-04S1), R01 DC016950, the Simons Center for the Social Brain and the Simons Foundation, and research funds from the Department of Brain and Cognitive Sciences and the McGovern Institute for Brain Research. The authors are grateful to all past EvLab members for help with data collection, especially Dima Ayyash, Jeanne Gallée, Alex Paunov, Nir Jacoby, Jayden Ziegler, Rachel Ryskin, Nafisa Syed, Saima Malik-Moraleda, Yotaro Sueoka, Yev Diachek, Elinor Amit, Tariq Cannonier, Jenn Hu, and Meilin Zhan. EF is also grateful to Nancy Kanwisher for mentorship and support, which laid a foundation for this line of work, and to Ted Gibson for the support over the years.

## Supplemental Materials

**SI-1:**
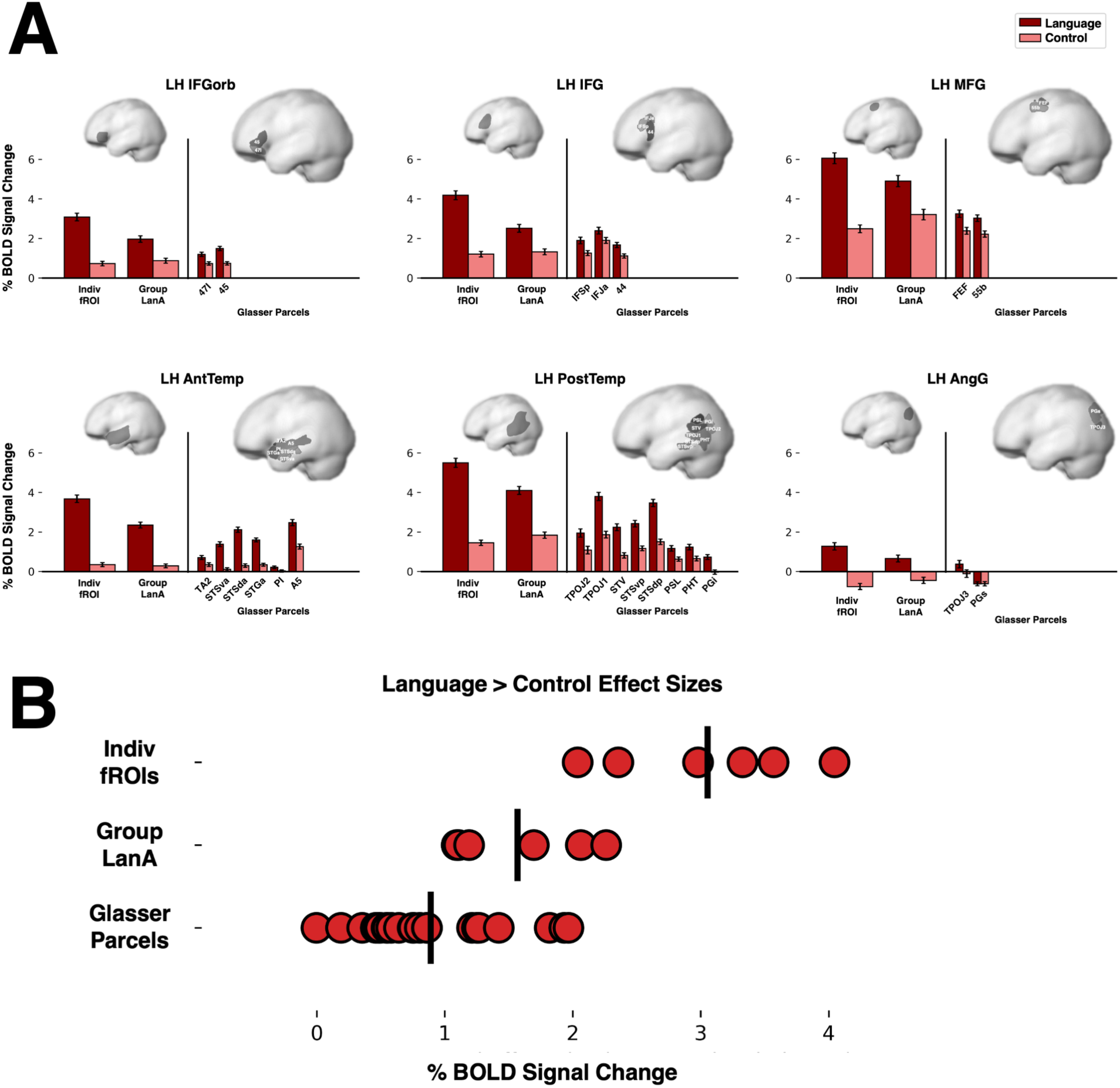
Response magnitudes for the language and control conditions and the size of the *language > control* effect for three sets of regions of interest (ROIs) in each of the six language parcels (shown on the left, smaller brain in each panel; see <u>Neural Markers</u>) for a subset of 403 participants: i) individual functional ROIs (Indiv fROIs), defined as described in <u>Neural Markers</u>; ii) ROIs based on LanA, where within each language parcel 10% of voxels with the highest overlap values in LanA are selected based on an independent set of participants (Group LanA; see below for details), and iii) ROIs based on the multi-modal data in the Human Connectome Project created by Glasser et al. (2016) (Glasser Parcels; shown on the right, larger brain in each panel). For the latter, we selected a subset of the parcels (n=23) that had at least 25% voxel overlap with one of the language parcels. Between 2 and 8 parcels overlapped with each of the six language parcels. *A)*. Magnitude of the *language* and *control* effects relative to the fixation baseline. Error bars reflect standard error of the mean (SEM). *B)*. Size of the *language > control* effect. For each set of ROIs, each dot corresponds to a language parcel (for Indiv fROIs and Group LanA) or to a Glasser parcel. Vertical black lines mark the average for that set of ROIs. As the figure clearly shows, individually defined fROIs fare best in terms of accurate estimates of effect sizes (and thus remain the gold standard), followed by the Group LanA ROIs, followed by the Glasser parcels. A few of the Glasser parcels in the temporal cortex perform comparably to the Group LanA ROIs, but in the frontal lobe all Glasser parcels grossly underestimate effect sizes (see Fedorenko &Blank, 2020 for a discussion of why frontal group-level ROIs are doomed to fail). Details on the definition of Group LanA ROIs: From the total set of 803 participants for whom we have two or more runs of a language localizer, we took a random sample of 400 participants. From those 400 participants, we generated a probabilistic volumetric atlas exactly as described in <u>Atlas creation</u>. For the remaining 403 left-out participants we defined ROIs as follows: within each language parcel, we select—based on the atlas created from the 400 independent participants—the 10% of voxels that have the highest overlap values, i.e., the voxels that are most likely to be language voxels. Critically, unlike for the individual fROIs, the same exact set of voxels is used in every participant, similar to anatomical or multi-modal group ROIs, like the Glasser parcels.

**SI-2:**
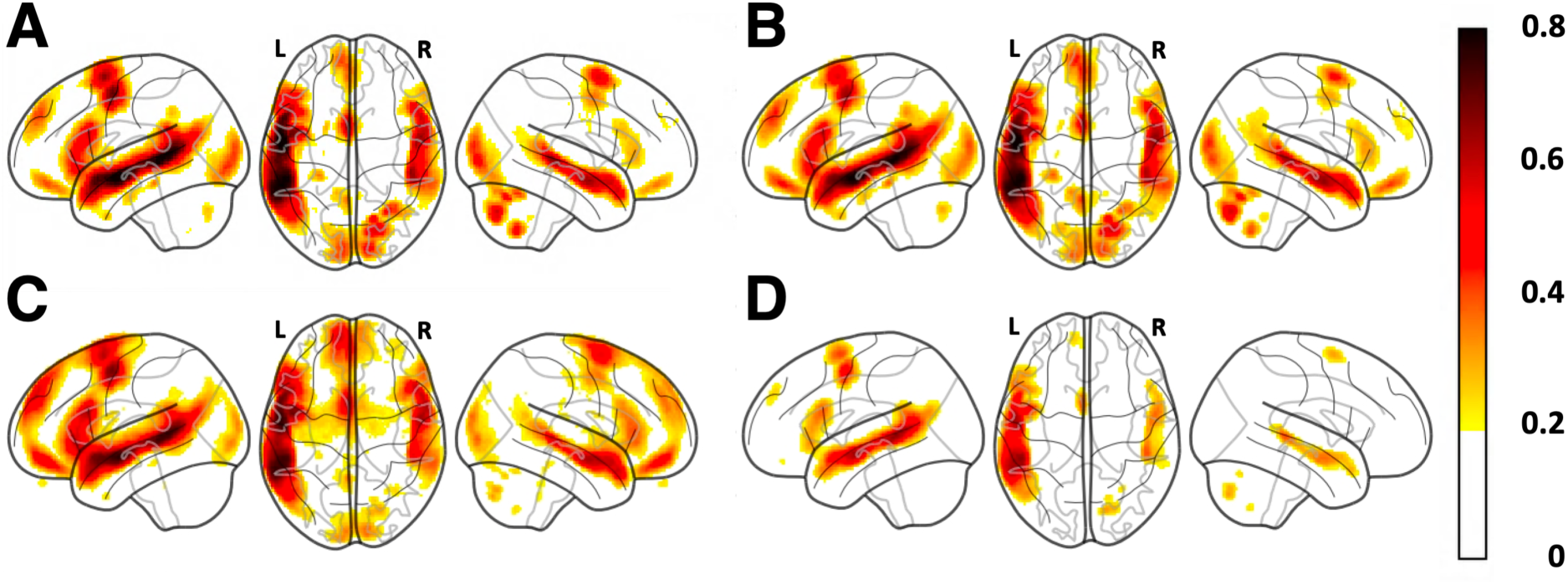
The robustness of the atlas to variation in data selection and other aspects of the procedure. Each alternative version of the atlas was compared to the original (top 10% of *language > control* voxels selected based on the *t-*maps across all 806 participants) by evaluating a spatial Pearson correlation over the set of all voxels in the atlas. *A)*. Atlas generated using only *Localizer Version A* (n=624 participants). Comparison to LanA: *r=*0.996. *B)*. Atlas generated using only native English speakers (n=629 participants). Comparison to LanA: *r=*0.998. *C)*. Atlas generated by selecting the top 10% of voxels from the contrast (*con*) maps rather than the variance-normalized *spmT* maps. Comparison to LanA: *r=*0.934. *D)*. Atlas generated using a fixed *t*-value threshold of 3.09 (∼p<0.001). Comparison to LanA: *r=*0.942 (although the overlap values were generally lower: maximum 0.66 in the LH (cf. 0.82 in the original atlas) and maximum 0.40 in the RH (cf. 0.64)).

**SI-3:**
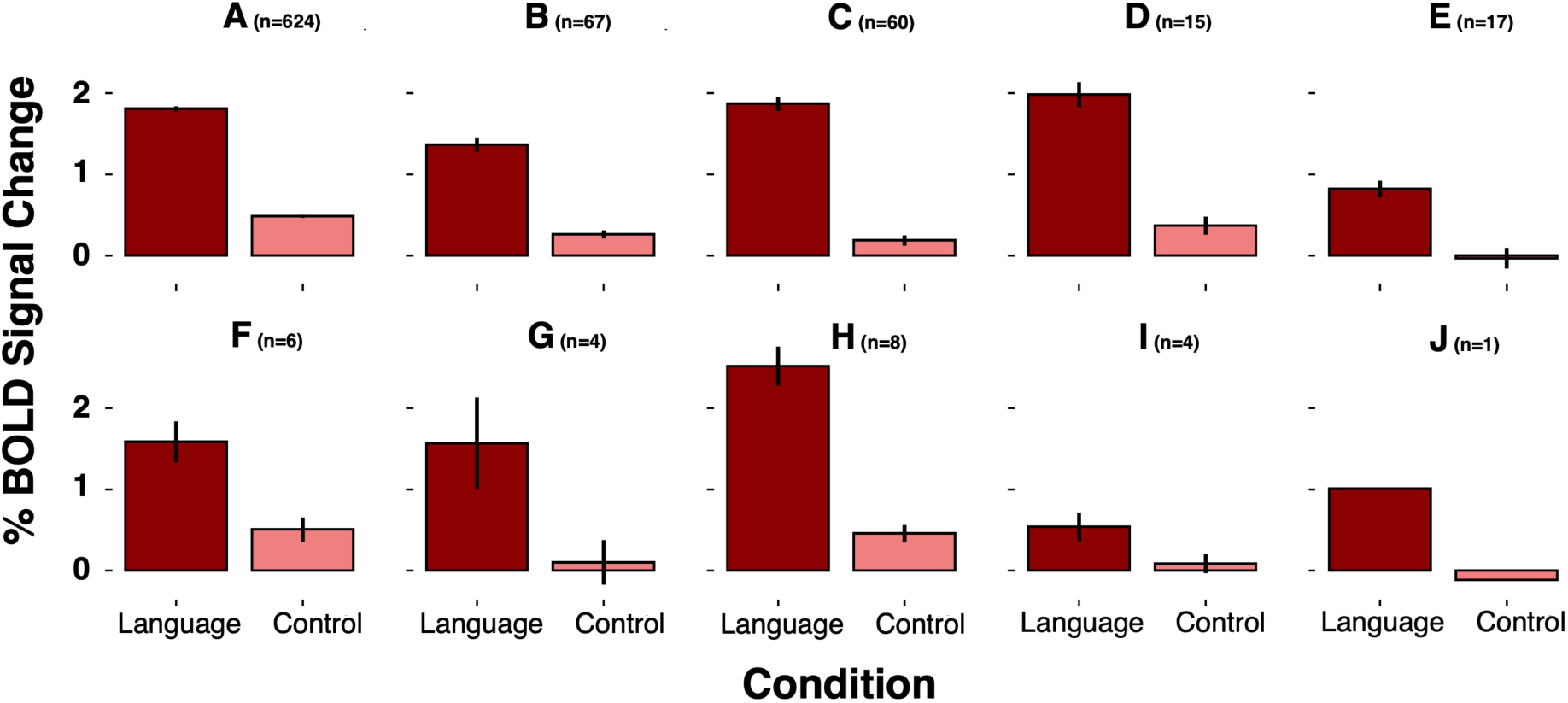
All 10 localizer versions (Table 2) show strong *language > control* effects. Error bars reflect standard error of the mean (SEM). For this analysis, we included 804 participants for whom we have at least 2 runs (necessary for across-runs cross-validation, as described in <u>Neural Markers</u>). For each participant, we calculated the average magnitudes across the 6 fROIs of the language-dominant hemisphere, defined as the hemisphere with the larger number of significant voxels (at the p<0.001, uncorrected threshold) within the union of the language parcels (see <u>Neural Markers</u>). Across all localizer versions, we see a strong *language > control* effect (in line with past work: e.g., Fedorenko et al., 2010; Scott et al., 2017).

